# SpaceWalker: Interactive Gradient Exploration for Spatial Transcriptomics Data

**DOI:** 10.1101/2023.03.20.532934

**Authors:** Chang Li, Julian Thijssen, Tamim Abdelaal, Thomas Höllt, Boudewijn Lelieveldt

## Abstract

Spatial transcriptomics (ST) enables profiling the expression of hundreds of genes in tissue sections, down to the level of single cells in their tissue environment. The gradient structure of ST data is particularly interesting for tissue biology, since spatial gene expression gradients often represent tissue compartment edges, whereas in the single-cell transcriptomic domain, gene expression gradients may represent cell type differences and smooth phenotypic transitions. Various computational approaches have been developed to extract information from either the spatial domain or gene expression domain individually. However, integrative biological interpretation of expression gradients in single cell and ST data spaces remains challenging. Many prior spatial transcriptomics analysis pipelines are script-based, lack interactive exploration facilities, and do not have specific facilities for automatic identification of localized expression gradients. Here, we present SpaceWalker, a visual analytics tool for exploring the local gradient structure of ST data. The user is guided by the local intrinsic dimensionality of the high-dimensional data to define seed locations, from which a flood-fill algorithm approximates k-nearest neighbor subgraph topology on the fly. In several use cases, we demonstrate that the spatial projection of these local subgraphs highlights tissue architectural features, and that interactive retrieval of gene expression gradients in the spatial and transcriptomic domains confirms known biology, and provides additional insights into the tissue architecture. We also show that SpaceWalker generalizes to several different ST protocols, and scales well to large, multi-slice, whole-brain ST data, while maintaining real-time interaction performance.

## 1. Introduction

Spatial transcriptomics (ST) enables profiling the expression of hundreds of genes at the level of single cells in tissue sections, thus providing new opportunities to understand tissue biology by combining single-cell gene expression profiles and their corresponding spatial context in the tissue1. For single cells, these High-Dimensional (HD) gene expression profiles enable detailed characterization of cell types, cell states and cell maturation^2^. Data visualization methods, and specifically Dimensionality Reduction (DR) techniques are extensively used to help understand complex HD data by giving a meaningful Low-Dimensional (LD) representation of these HD spaces. This typically consists of generating and visualizing a LD map in which distances, similarities or neighborhood relations from the HD data space are preserved in the LD space^3^. DR embeddings give insight into the structure of the data, showing clusters and generic trends in the data. Besides analysis of which cell types are present in a sample, trajectory inference approaches aim to computationally derive transitions between cell types and states and order cells along a trajectory topology. This allows investigating cellular dynamics, where expression gradients in one or more genes encode biologically relevant state transitions and cell variability^4^. DR techniques are typically applied to reduce data complexity, and algorithms like k-Nearest Neighbor (kNN) graph construction and clustering techniques are used to extract the topological structure of the data2.

As single-cell RNA sequencing (scRNA-seq) does not support characterizing the spatial organization and interactions between cells, ST adds a spatial dimension to single-cell analysis, enabling the study of cell-cell interactions, tissue architecture, and coupled cell development and migration trajectories in the tissue5. Understanding the structure of the spatial map or non-linear DR embeddings often requires localized methods to visualize the features driving the map structure at that particular location6,7. Coloring the cells in these 2D maps with gene expression values is commonly used in single-cell analysis to visualize the underlying expression patterns, but genes of interest have to be manually selected and it is not possible to reveal how cells are connected in HD space.

Various computational approaches have been developed to extract information from either the spatial domain or the gene expression domain. For instance, Hidden Markov random fields were used to integrate scRNA-seq data and spatial neighborhood information8-10. In addition, several methods exist for spatial gene expression analysis. SpatialDE11 applied a Gaussian process regression to identify genes with correlated expression levels in the spatial domain. Trendsceek^12^ detected spatial dependency of gene expressions using a marked point process. Although these approaches enable the identification of spatial gene expression variation, localized gene expression analysis methods are lacking. Furthermore, these computational strategies are script-based and lack interactive data exploration facilities with a direct feedback loop to the user.

Explorative analysis and visualization of ST data allows for generating hypotheses on tissue biology. Integrative toolboxes such as Giotto13 and Squidpy14 allow researchers to interactively visualize and analyze spatial data. Cytosplore15 is software that uses t-SNE16 projection as the main view for real-time interactive visual exploration in the single-cell analysis domain. In addition to efficiently processing large-scale mass cytometry data, it also provides clustering techniques and supports multiple linked views between the map structure and the feature space. Based on Cytosplore, Cytosplore Transcriptomics^17^ provides interactive analysis for scRNA-seq data. CELLxGENE18 is a web-based interface aimed at interactive exploration of HD single-cell datasets. It enables collaborative analysis between experimentalists and bioinformaticians. Despite the active development of interactive visualization tools, interactive biological interpretation of single-cell HD and spatial data remains challenging. The relationship between HD single-cell data and 2D maps has not been fully explored.

In this work, we present SpaceWalker, a visual analytics tool for exploring the gradient structure of ST data. Specifically, we focus on interactive exploration of localized gene expression gradients: these are particularly interesting for tissue biology, since spatial gene expression gradients often represent tissue compartment edges, whereas in the HD single-cell transcriptomic domain, they represent cell type differences and smooth phenotypic transitions between them. In SpaceWalker, the user is guided by the local intrinsic dimensionality of the HD data to interactively pick seed locations for a series of neighborhood searches on the kNN graph. The results of these searches, i.e., localized HD neighborhood “flood-fills”, are then projected onto the 2D spatial map in real time, revealing the spatial topology of the HD kNN graph; These localized HD topology approximations then serve as input for gradient detection filters, which prioritize genes with a localized (spatial or HD) expression pattern. The genes that exhibit a localized spatial or HD gradient are visualized in real-time in the spatial domain, along with a number of options to enable user-tailored data exploration paths. This offers the user a real-time querying of the gradient structure of ST data. In several use cases, we demonstrate that the spatial projection of these local kNN subgraphs highlights tissue architectural features, and that interactive retrieval of gene expression gradients in the spatial and transcriptomic domains confirms known biology, and provides additional insights in tissue architecture. We also show that SpaceWalker generalizes to several different ST protocols, and scales well to large, multi-slice, whole-brain ST data, while maintaining real-time interaction performance.

## 2. Results

### SpaceWalker methodology

#### Overview

The goal of SpaceWalker is to provide the user with an interactive interface for exploring localized expression gradients in ST datasets. By offering highly responsive global and local linked views of cells and genes, SpaceWalker aims to help users identify tissue architecture as well as locally variable genes, and to gain insights that would be difficult to uncover using script-based methods. An overview of the proposed methodology is shown in Figure 1. The input of SpaceWalker is a cell-by-gene expression matrix, with spatial coordinates assigned to the single-cell gene expression vector19. The idea of SpaceWalker is to guide the user exploration by a global overview of local intrinsic dimensionality at each cell location (Figure 1(a)). From a user-selected seed cell, the projection of the local HD structure is estimated and highlighted in the spatial map in real-time, i.e., cells with similar transcriptomic profiles to the seed cell are presented to the user (Figure 1(b)). Genes with spatially localized expression peaks in the area of the selected cell are detected using a spatial filter kernel (Figure 1(e)) that ranks genes by localized expression variation. Alternatively, a filter for localized expression peaks in the HD space (Figure 1(f)) can be selected. These filters effectively serve as real-time gene image retrieval based on localized expression variability. Since multiple genes can exhibit similar filter responses, we also provide a line chart of all genes, showing sorted gene expressions in the local neighborhood (Figure 1(c)). This offers a complementary view to the filter gene rankings. The user can click on a gene of interest in the chart, and inspect the gene expression of the chosen gene in a separate gene view (Figure 1(d)). This allows them to not only inspect the genes automatically selected by the filter, but to also manually explore genes with interesting expression values in the local neighborhood.

**Figure 1:**
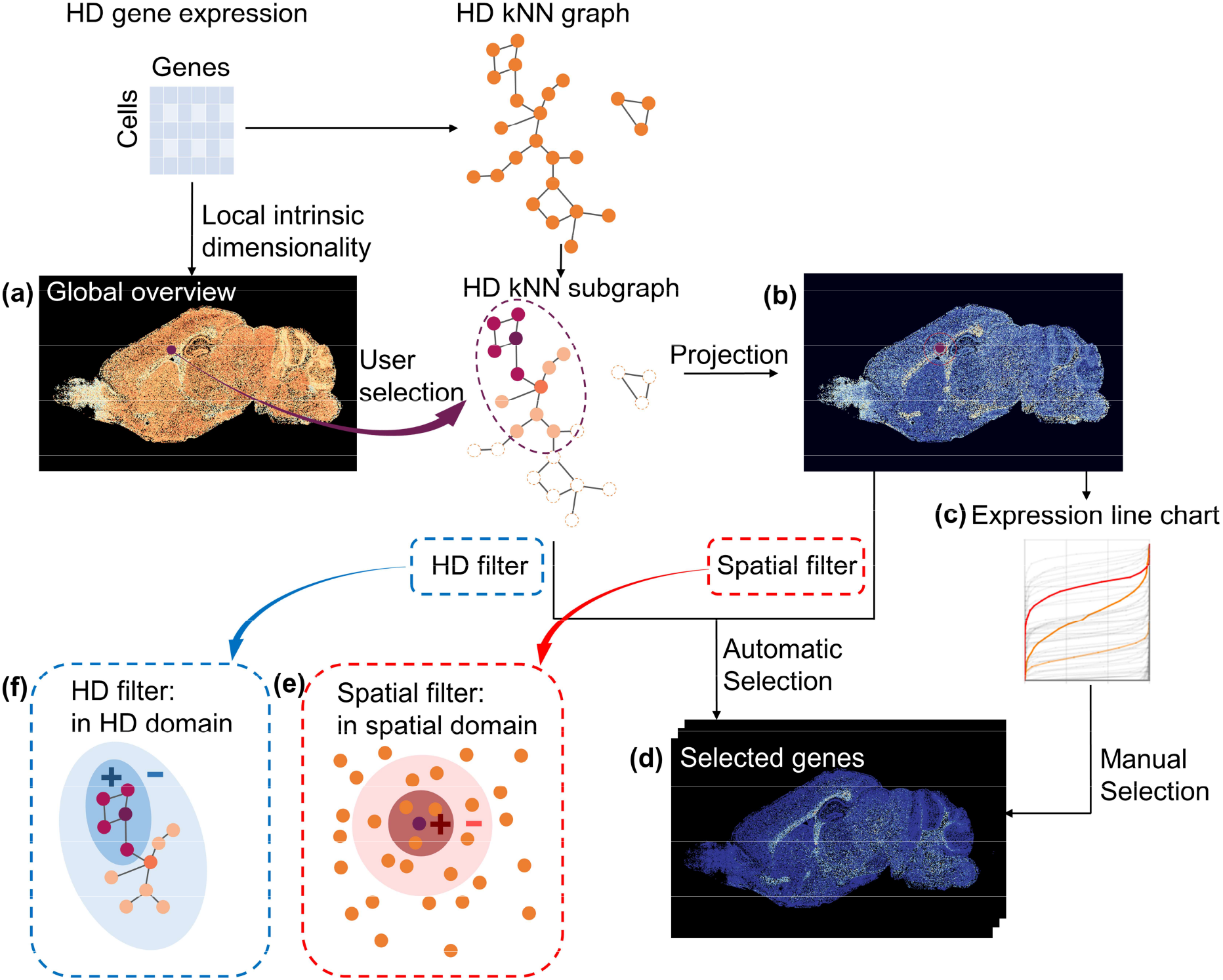
Overview of SpaceWalker. The user interacts with a 2D scatter plot (a), where the spatial map is color-coded with the local intrinsic dimensionality of the HD data to highlight areas of potential interest. A flood-fill algorithm approximates the local topology of the HD space at the selected cell node, and projects it back on the spatial map (b), displaying the HD structure and cell similarities. Genes with localized expression peaks are ranked automatically via a spatial (e) or HD filter kernel (f), all genes are presented in a line chart (c) with their sorted expression values in the local neighborhood. The user can inspect the gene expression (d) of top ranked genes (automatic selection) and can also manually select different genes of interest from the gene expression line chart (manual selection). Each line in the chart (c) represents a gene, and genes ranked the highest by the filter are highlighted in orange and the manually selected gene is highlighted in red.

#### kNN subgraph projection

First, a kNN graph is constructed from the whole gene expression profiles of individual cells in the ST data. Multiple distance metrics can be used when constructing the kNN graph, including Manhattan distance, Euclidean distance, cosine distance, as well as secondary similarity measures20 to improve quality on higher-dimensional datasets. To ensure scalability towards large datasets and optimal computational speed, we deploy the GPU-based kNN library FAISS^21^, which provides efficient similarity searching for the construction of the kNN graph. To enable real-time extraction of local graph structure, we traverse the kNN graph starting from the direct, or 1st order neighborhood to the n-th order neighborhood, where n is a user-defined number of indirections, or steps along the graph. When the user selects a seed cell in the spatial map, we construct its local HD neighborhood subgraph by using this computed kNN graph. Starting from the selected cell, its direct neighbors, as dictated by the kNN graph, are added to the subgraph. Then, for n steps, the direct neighbors of the cells added in the previous step are added to the subgraph. Neighbors that are already present in the current subgraph are not added. This process can be seen as a flood-fill from the seed cell over its HD topology, and will be referred to as flood-fill in the rest of this work. The term “flooded cells” refers to the cells that have been added to the subgraph in an n-th order neighborhood manner for n steps.

Importantly, this process differs from simply computing the seed cell’s nearest neighbors with a higher k value, as each wave of the flood is restricted to a local neighborhood graph. This makes it improbable that a wave spreads to dissimilar cells as those are unlikely to be part of the local neighborhood. On the other hand, computing the cell’s nearest neighbors with a large k value can easily cross to dissimilar clusters in the HD space if the size of k exceeds that of the local cluster. Therefore, the flood-fill algorithm more closely approximates the neighborhood of similar cells and extinguishes itself when it reaches the boundaries of that neighborhood.

#### Whole-slide visualization of local HD intrinsic dimensionality

The visual exploration in SpaceWalker is guided by features that express the local variance of the HD gene expression space, as a proxy for biological variability (Figure 1(a)). To this end, the local intrinsic dimensionality is computed for a neighborhood around each data point in HD gene expression space and used to color the spatial map. This provides a global overview of region boundaries to guide the spatial exploration by transcriptomic variability. Intrinsic dimensionality for the k nearest neighbors of each cell is computed by applying PCA22, and recording the number of principal components required to model 85% of the total local variance. The total variance explained by these intrinsic dimensions is the locally explained variance in the k nearest neighbors.

In addition to features expressing local variance, we also provide exploration guidance in the form of a metric for local density of the HD space. The flood-fill algorithm used to identify the nearest neighbors of a seed cell is controlled by a fixed number of steps, resulting in varying flood-fill sizes at each seed cell location. The flood-fill size reflects the HD density of the dataset, with a larger neighborhood indicating a sparser HD structure and a smaller neighborhood indicating a denser structure.

#### Localized gene filtering and selection

ST data is often used to explore boundaries between transcriptomically distinct tissue regions and cell mixtures, differentiation trajectories or cell migration paths. Such patterns are typically characterized by localized changes in the expression of one or more genes. To enable the study of such localized gene expression gradients, we developed an exploration mode that enables the user to interact on the 2D map, while at the same time genes with significant localized expression patterns are ranked by two filters: 1) a spatial peak filter (Figure 1(e)), representing differential expression between the average gene expression vectors of two 2D circular neighborhoods with different radii: such filters contrasting two different spatial neighborhood sizes are common in classical image processing23; 2) a HD filter (Figure 1(f)), contrasting the average gene expression vectors within two flooded HD neighborhoods of different number of steps. We opted for peak filtering by computing differential expression between a small and a large (spatial or HD) neighborhood due to its low computational complexity and rotational invariance in the spatial domain. Additionally, peak filtering has the capacity to detect both peaks and gradients with the same filter, since the gradient is often near the peak, reflecting the transition from high to low values. The filter neighborhood sizes can be modified through interactive sliders on the user interface, where the ratio between neighborhoods shifts the filter properties between peaks and gradients.

The aforementioned filters enable an automated detection of genes with localized expression patterns by sorting genes according to filter response. The top-ranked genes are presented in linked panels for their spatial expression patterns, and can be inspected by clicking on the gene panel of interest (Supplementary Movie 1 and 2). However, co-expressing genes may have slightly lower filter ranks, therefore we enable the user to interactively deviate from the automated gene selection based on ranks. In a line chart, we plot a series of lines, where each line represents the sorted (low-to-high) gene expression of that particular gene over the flooded cells (Figure 1(c)). Genes with a high-ranked filter response are highlighted on the chart. While the user explores different cell locations on the map, gene expression values of the flooded cells are updated in the line chart, providing an overview of local gene expression patterns. The user can interactively select a line in this chart to see the spatial gene expression map of the corresponding gene. This enables the user to investigate other genes of interest based on their gene expression profiles or prior knowledge. For example, a gene showing high expression values throughout all of the cells within the flood-fill might indicate a large contribution to the flooded neighborhood. Alternatively, a gene showing a gradual change in gene expression could point to cell maturation trajectories.

Finally, we implemented the option to apply the spatial filter only on the trancriptomically similar flooded cells, or on cell subtypes selected from a cell-type taxonomy. This enables the exploration of spatial expression gradients within isolated cell subtypes (e.g. glutamatergic neurons) without mixing of cell types in the gene filter region of interest.

#### Coloring flood-fills by localized features

Flooded cells can be colored with different features, i.e. foreground cells are highlighted, and colored with a different feature than the background cells (Figure 6 (d1) (d3)). These localized color-coding options assist the user in exploring the local subspace properties in the context of HD geodesic distances on the kNN graph and expression gradients. The user can then interact with the map and explore local data properties by selecting one of the following flood-fill color-codings:

- Geodesic distance approximation on the kNN graph: Flooded cells are color-coded by their flood-fill step index to indicate their proximity to the seed cell in HD space, hence reflecting local HD kNN graph geometry. The user can interactively define the number of flood-fill steps to inspect the HD topology.
- Gene expression: Flooded cells are colored with the expression of the top ranked gene, and all other cells with a constant background color. The user can also manually select other genes of interest from the gene expression line chart for coloring the flooded cells. The localized view of gene expression enables the user to visually evaluate the spatial geometry of the flood-fill in combination with the gene expression pattern.

### Real-time projection of kNN subgraph structure in the spatial domain reveals tissue-architectural features

First, we investigated whether local tissue architectural features are reflected and preserved by projecting the localized HD neighborhoods on the spatial data. To this end, we used two publicly available ST datasets of the mouse visual cortex that were acquired in the context of the SpaceTx consortium24. We selected these datasets (smFISH and MERFISH) because of the abundance of literature on the laminar spatial architecture of the mouse visual cortex^25^. The SpaceTx datasets contain manual annotation of the cortical layers, which enables investigating the robustness of SpaceWalker against ST protocols applied to the same tissue sample (smFISH and MERFISH).

To quantitatively assess correspondence of the local neighborhood geometry with local tissue architecture, we defined a reference standard based on the known laminar structure of the mouse visual cortex. We defined the *Layer Percentage (LP)* of flooded cells that have the same layer annotation as the seed cell: this assumes that transcriptomically similar cells are organized in spatial layers, which is the case for excitatory neurons in the visual cortex. For any selected cell, the projection of the local HD neighborhood reveals the spatial organization of cells with similar expression profiles (Supplementary Movie 1). Flooded neighborhood projections for all cells in the tissue clearly reveal that the flood-fill neighborhood searching recovers the laminar tissue architecture (Figure 2). Cells that are not located near the layer boundary on the spatial map show a high *LP* value, demonstrating that the local HD neighborhood projection accurately reveals the known laminar architecture of cells with similar expression profiles. Cells located near the boundary of the spatial layer are less likely to be in the same layer as the manual seed cell annotation, and thus show a lower *LP* value. Another explanation for the lower *LP* values could be due to the fact that transcriptomically similar cells may appear in multiple layers^25^. This cell type cross-talk across cortical layer boundaries explains why flood-fill neighborhood searching seeded close to the manually annotated layer boundaries may span across the manually annotated layer boundaries. However, the results obtained from the independently acquired smFISH and MERFISH datasets exhibit nearly identical layer patterning in Figure 2, indicating robustness of Spacewalker against ST protocol, and reproducible tissue architecture recovery across datasets.

**Figure 2:**
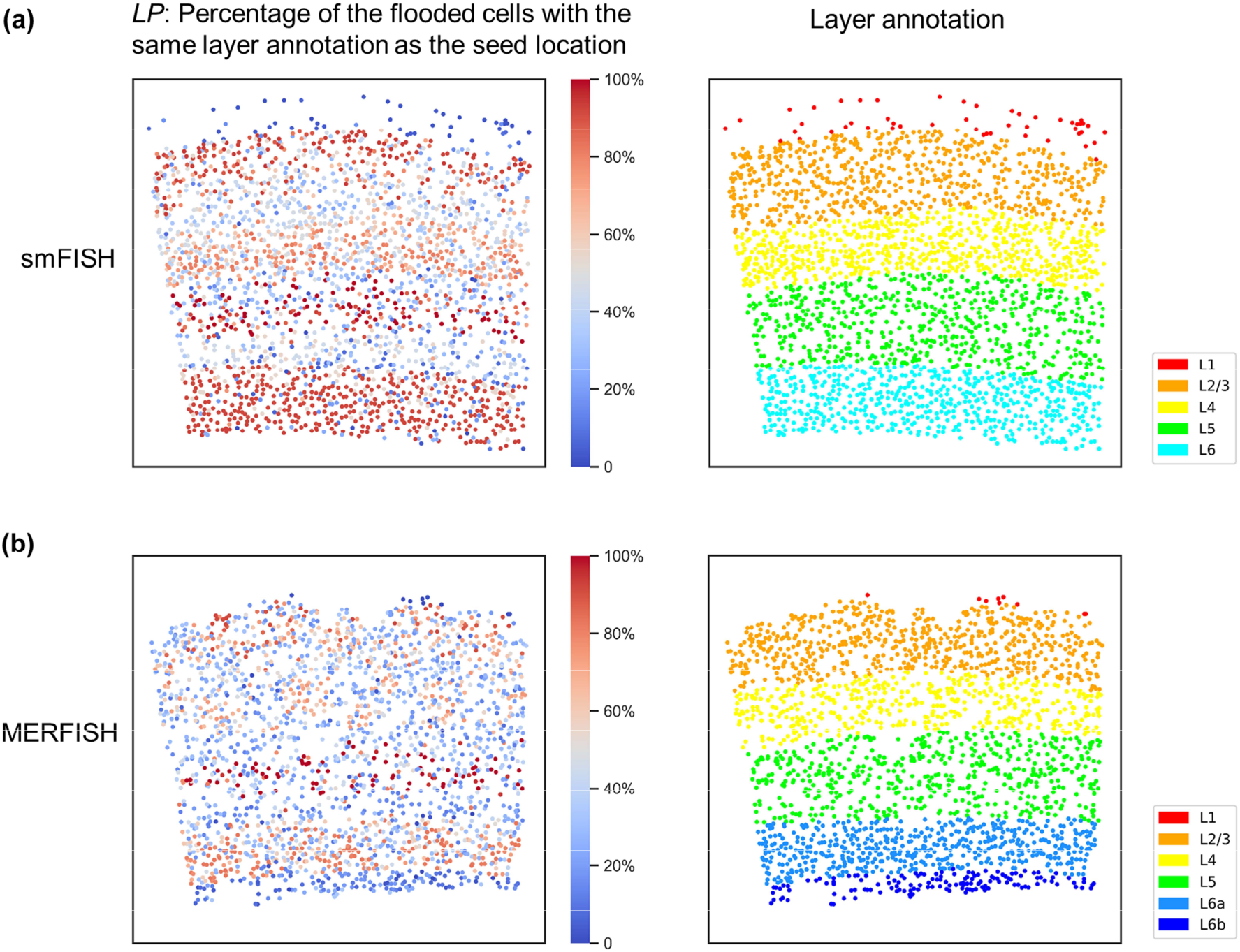
Spatial tissue maps color-coded by *LP*: the percentage of flooded cells with the same layer annotation as the seed cell of smFISH (a) and MERFISH (b) datasets and their manual layer annotation.

To investigate whether SpaceWalker neighborhoods also reflect more complex spatial geometries accurately, we applied SpaceWalker to HybISS data of the developing mouse brain26. When projecting the local HD neighborhood back to the 2D spatial map, the cells are colored by their flood-fill step index, which represents a measure for the HD geodesic distance to the seed point. The neighborhood mapping presents similar patterns as the gold standard region labels that were imputed from single-cell transcriptomics data26 (Figure 3(a)). For example, HD neighborhood projections highlighted the structure of hindbrain floorplate and mesenchyme, even if the two starting cells of the flood-fill algorithm were located next to each other in the spatial map (Figure 3(b, c)). HD neighborhood projection also highlighted the similarity between ventral hindbrain cells that are not located close by in the spatial map (Figure 3(d)).

**Figure 3:**
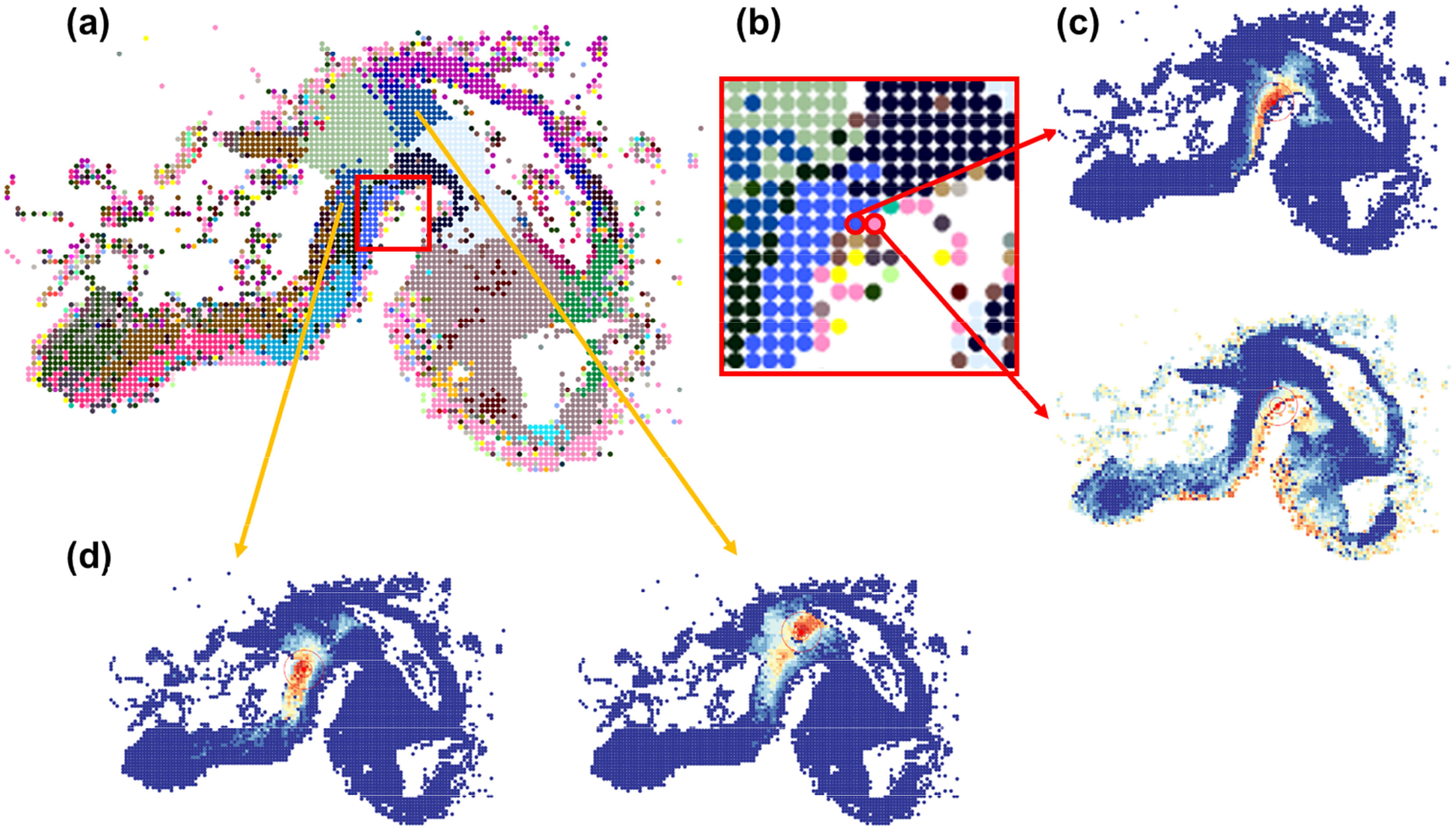
HD neighborhood projections by flood-filling agree with subclass annotation in HybISS data. (a) Spatial map color-coded with cell subclass annotation26. (b) Magnified view of (a) showing two neighboring cells in the spatial map belonging to hindbrain floorplate and mesenchyme respectively. (c) HD neighborhood projections seeded from the two marked cells in (b), highlighting distinct spatial patterns that coincide with the region labels of hindbrain floorplate and mesenchyme. (d) HD neighborhood projections starting from two ventral hindbrain cells that are located in two different regions in the spatial map, highlighting the transcriptomic similarity between the two different ventral hindbrain regions.

### Real-time gene filtering reveals localized gene expression peaks and gradients in the spatial domain and HD domain

To quantitatively evaluate to what extent the top-ranked genes detected by the spatial and HD filters corresponded to known layer-specific marker genes in the mouse visual cortex25, we calculated a frequency count for filtered genes by counting how often genes end up in the top two ranks at different seed cell locations per cortical layer.

Apart from detection capacity for known marker genes, we aimed to assess whether SpaceWalker can also detect previously unreported genes that exhibit clear laminar expression patterns. As such, we defined a quantitative metric for the layer-specificity of all genes by computing the correlation between gene expression images and the manual layer annotations. For each cortical layer, a layer mask was constructed based on the manual layer annotations, with cells within a specific layer encoded as 1 and all other cells encoded as 0. The correlation scores between all genes and the gold-standard layer masks are used as benchmark reference for the evaluation of filters. Finally, to assess robustness with respect to ST protocol, we computed both metrics for the smFISH as well as MERFISH data from the SpaceTx consortium.

Figure 4 shows a side-by-side comparison of the heatmaps of frequency counts of filtered genes (applying the spatial filter) and their correlation scores of smFISH data (results of MERFISH data are given in Supplementary Figure 1). All layer-specific marker genes have a high correlation score with their corresponding layer annotation, proving that the correlation metric identifies genes with contrasting patterns within a specific layer. Known marker genes as reported25 are presented in the top of the heatmaps, and the other genes are grouped based on layer and correlation score. Genes with a high frequency count in a specific layer also exhibit a high correlation score with the manually annotated layer masks, demonstrating the ability of the filters to capture local spatial variability. Some genes with a high ranking in our filters are not mentioned in the literature as known layer markers, even though they were highly correlated with the simulated layer masks. Visual inspection of these genes revealed clear laminar expression patterns (Figure 5).

**Figure 4:**
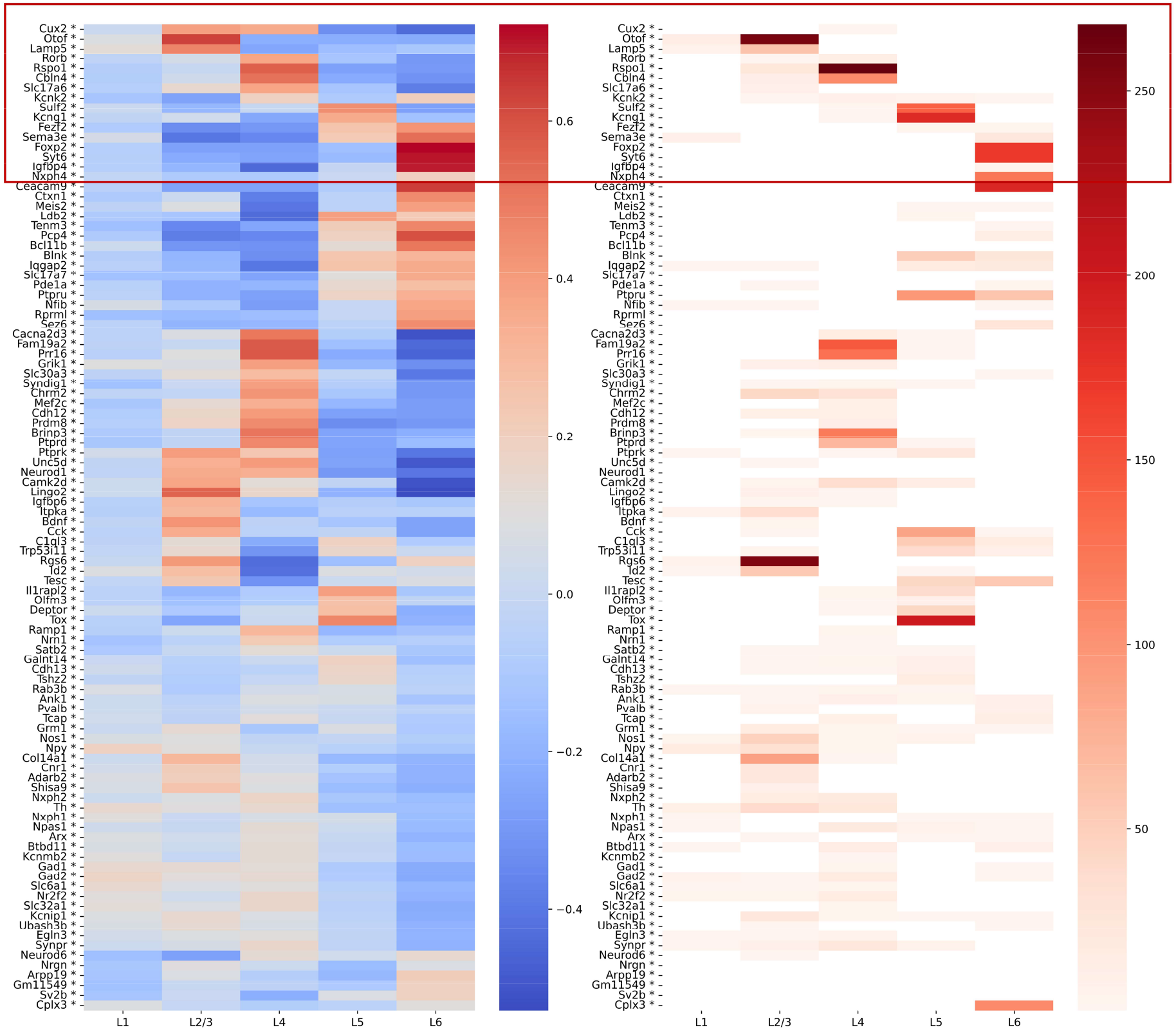
Heatmaps comparing the frequency counts by the spatial filter (right) with the correlation scores between genes and layer annotation masks (left) of smFISH dataset. Known marker genes as reported25 are highlighted in the red box. Genes with a high correlation with a layer mask were also often ranked in the top two by the spatial filter. Results of MERFISH dataset are given in Supplementary Figure 1. Results for the HD filter showed similar correspondence between correlation scores and frequency counts, and are given in Supplementary Figure 2, 3.

**Figure 5:**
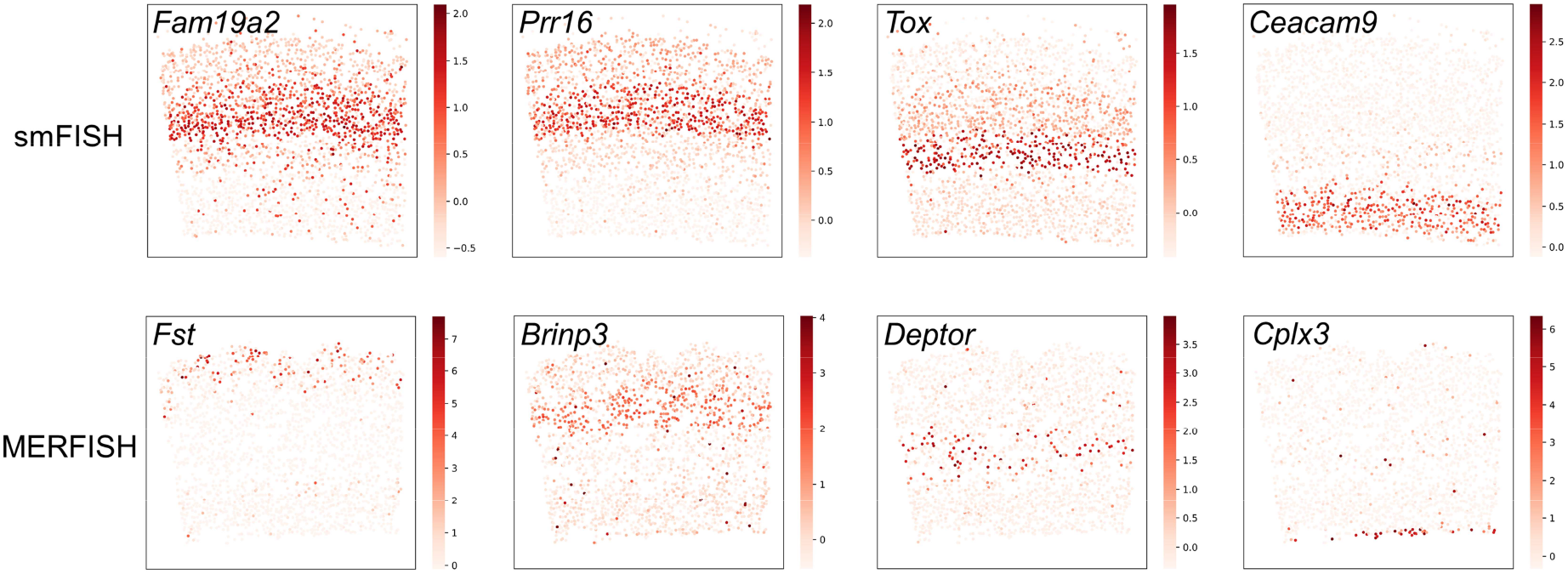
Examples of genes that are not mentioned in the literature as known layer markers, but frequently ranked as top two by the spatial filter.

It is important to note that the filter results do not perfectly align with layer-specific analysis results, as the spatial filter is designed to identify genes with local variability within the filter kernel, while layer-specific analysis is contrasting the expression vectors inside and outside of a layer. The filter size used in the comparison with layer-specific analysis is approximately equal to the spatial size of the layer, providing a basis for comparison. However, the user can adjust the filter size based on their interests to define the locality of the gene expression patterns.

As the filters only take cells within the filter kernel size into account, it is possible that layer-structured genes are detected that are also highly expressed in other locations (Supplementary Figure 4). It can also be seen in the heatmaps in Figure 4 and Supplementary Figure 1 that a gene can be ranked as top two in multiple layers as it has high correlation scores with multiple layers.

### Color-coding the spatial map with localized HD features provides visual cues for guidance of exploration

Figures 6(a) and 6(b) give examples of spatial maps in the SpaceTx datasets, color-coded by HD intrinsic dimensionality and flood-fill size. The laminar structure clearly emerges from the intrinsic-dimensionality coloring (consistent in both smFISH and MERFISH datasets) indicating that local intrinsic dimensionality provides visual cues for spatial tissue partitioning. The local intrinsic dimensionality of L6 is found to be higher than other layers, while the locally explained variance of L6 is lower. This suggests that the cells in L6 are less homogeneous than cells in other layers. L5 has a relatively dense HD structure compared to other layers, where the flood-fill is confined within a dense cluster. In contrast, the flood-fill progresses further in layers with relatively sparse HD structures, leading to a larger neighborhood size. It can be confirmed by t-SNE maps also color-coded by flood-fill size, where cells in L5 are divided into multiple smaller clusters while the cells in the other layers form distinct and well-separated clusters (Figure 6(b)).

**Figure 6:**
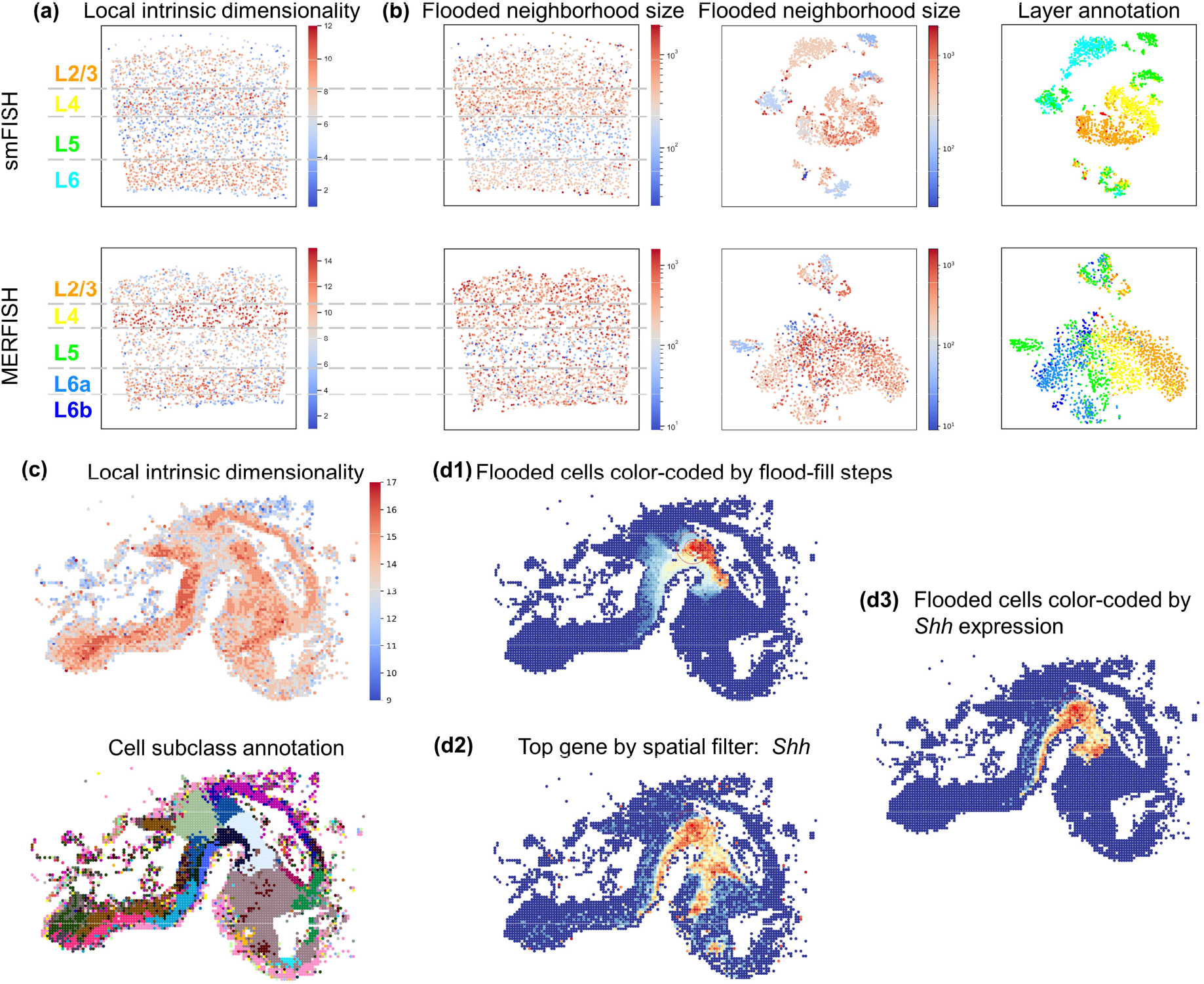
(a) Spatial maps color-coded by local intrinsic dimensionality of smFISH and MERFISH datasets. (b) Spatial maps color-coded by flooded neighborhood size (left) and t-SNE maps color-coded by flooded neighborhood size (middle) and layer annotation (right) of smFISH and MERFISH datasets. (c) Spatial maps of the HybISS dataset, color-coded by local intrinsic dimensionality and cell subclass annotations. (d) Color-coding of flooded HD neighbors by flood-fill index (d1) and *Shh* expression (d3), where *Shh* was ranked as the top gene by spatial filtering at the defined location. The entire spatial map colored by *Shh* expression is shown in (d2) as a reference.

Figure 6(c) shows how the intrinsic dimensionality can serve to guide the user to anatomically distinct regions in HybISS data, demonstrating that changes in local intrinsic dimensionality in many cases mirror transitions between cell subclasses. Color-coding of the flood-fill cells by flood-fill step index (Figure 6(d1)) reveals the local HD topology on the spatial map, whereas color-coding by gene expression of the top ranked *Shh* gene (Figure 6(d3)) reveals spatially localized gene expression.

### SpaceWalker scales to multiple whole mouse brain sections at real-time interaction speeds

In the above sections, we have used two SpaceTx datasets and a HybISS dataset to validate the functionality of SpaceWalker. The samples of mouse visual cortex in the SpaceTx datasets have a well-characterized tissue architecture and feature structure, demonstrating real-time performance on these datasets on consumer-grade computing platforms. These datasets are relatively small in scale (Table 1). Scalability in biological data analysis is considered of significant importance due to the increasing complexity of data14,18,19, as laboratory technologies continue to evolve. ST protocols such as EEL FISH19 now scale towards large Field of View (FoV) whole-organ tissue patches; Alternatively, through stitching of multiple patches, large tissue surface areas can now be compounded to a large FoV.

**Table 1:** Performance of SpaceWalker on different datasets. All experiments were conducted on a 6-core, 3.80GHz Intel processor with 128GB of RAM and an NVIDIA Quadro 2200 GPU.

|  | SpaceTx smFISH | SpaceTx MERFISH | HybISS | EEL FISH | Vizgen MERFISH (multi-slice) |
| --- | --- | --- | --- | --- | --- |
| Data dimension [cells,genes] | [2360, 314] | [2150, 314] | [4628, 117] | [127591, 440] | [161875, 483] |
| Pre-processing (s) | 1.56 | 1.53 | 0.89 | 8.76 | 19.68 |
| Memory usage (GB) | 1.04 | 1.02 | 1.19 | 1.75 | 2.36 |

To investigate whether SpaceWalker still facilitates real-time exploration at such large patches, we deployed it to a sagittal whole-brain slice acquired with EEL FISH19. This dataset consists of 120k cells and 440 genes and it takes 8.76 s to load in SpaceWalker, including data preparation, kNN computing, and initialization. The user can then start interacting smoothly with the interface based on real-time computation. An overview of computation times and memory footprints for all experiments in this paper are given in Table 1.

Next, we investigated whether SpaceWalker exploration of this large dataset maintained similar tissue architecture retrieval and feature detection performance as with the smaller datasets reported above. Figure 7 gives examples of local intrinsic dimensionality based on two different kNN graphs, local flood-fill reprojections and identification of genes with localized expression patterns detected by the spatial filter. Cells in the cerebellum, corpus callosum and olfactory bulb exhibit lower intrinsic dimensionality than other tissue areas, delineating anatomical region boundaries (Figure 7(b)). The local neighborhood projections (Figure 7(c)) highlighted the anatomical regions in EEL FISH and identified genes of which the spatial expression profile resembled the flood-fill geometry (Figure 7(d)). Also, several of the filter-ranked genes were detected that corresponded to genes reported^19^ (*Mbp, Plp1* in the corpus callosum; *Drd2* in the striatum; *KI* and *Otx2* in the choroid plexus; *Ramp3, Synpo2* in the thalamus; *Cbln1* and *Fam107a* in the cerebellum). These results demonstrate that SpaceWalker scales to large, whole-slide ST data while maintaining interactive speed (Supplementary Movie 2), and still retrieves genes that confirm known local tissue biology.

**Figure 7:**
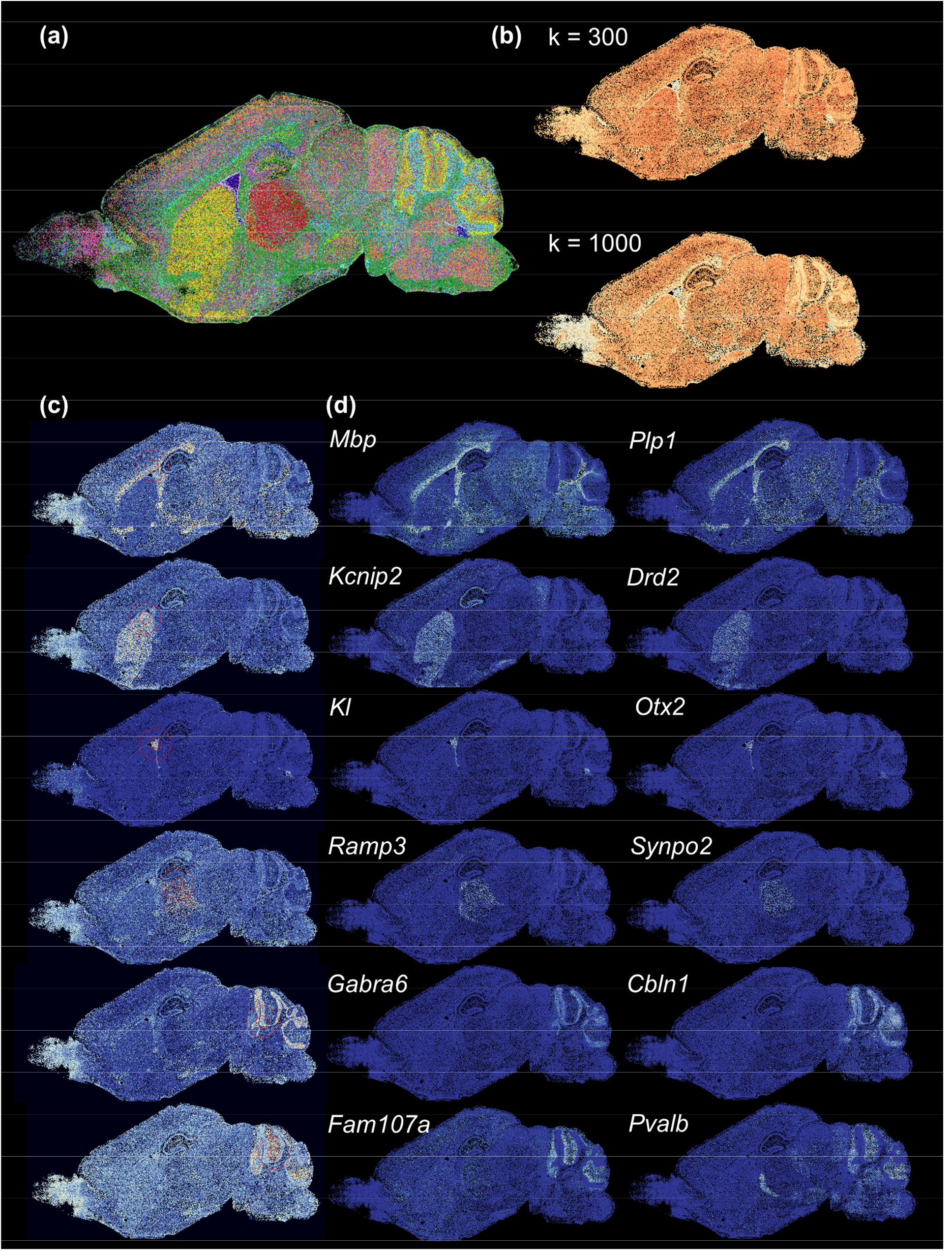
Exploration results of EEL FISH dataset revealing tissue structure and local gene patterns. (a) Reference annotation19, in which 30 selected genes highlight anatomical structures of a sagittal mouse brain section. (b) Spatial maps color-coded with local intrinsic dimensionality with k=300 and k=1000, indicating that local intrinsic dimensionality provides visual cues about region boundaries. (c) Examples of local neighborhood reprojections, flood-fill cells are colored by flood-fill step index from red to light color, and background cells with dark blue. (d) Top two genes selected by spatial filtering at the location of seed cell in (c), each highlighting spatial gene expression that is similar to the flood-fill geometry.

As we demonstrate above, SpaceWalker is capable of exploring large-scale ST datasets at interactive performance. SpaceWalker is not only limited to handling one single spatial map, it also supports multi-slice spatial map exploration by pooling data from multiple slices. With the recent emergence of whole-brain ST atlases^27^, exploring multiple slices at the same time provides additional information across the slices. The similar tissue regions across multiple slices are highlighted, as the kNN graph and flood-fill algorithm were based on pooled data from all slices. By visualizing and exploring multiple slices, it provides a possibility to explore the structure of the whole brain dataset as a proxy for a 3D representation.

To demonstrate a multi-slice use case scenario, we deployed two full coronal slices from Vizgen MERFISH Mouse Receptor Map dataset (Vizgen Data Release V1.0. May 2021^28^). Multi-slice exploration reveals relationships between slices. Related tissue structures are highlighted in both slices during exploration on one of the slices (Supplementary Figure 5). Genes that display a localized expression peak in both slices are ranked high by the filters. It is important to note that SpaceWalker is capable of handling datasets with more than two slices. SpaceWalker remains performant and fully interactive despite the scalability challenge that multi-slice data might present.

## 3. Discussion

Interpretation and analysis of ST datasets is often limited to script-based tooling with limited interactivity. However, hypothesis generation from complex biological data benefits from interactive, on-the-fly querying of the data, especially for ST data, since genes can exhibit different functions at different locations in the tissue. This also holds for localized spatial or transcriptomic expression gradients: spatial expression gradients are important indicators for tissue boundaries, migration trajectories and spatially evolving cell phenotypes, whereas transcriptomic gradients reflect cell states, differentiation and cell-type boundaries.

Here, we present SpaceWalker, an interactive visual analytics tool for exploration of large patches of ST data. SpaceWalker consists of two key innovations: an interactive, real-time flood-fill and spatial projection of the local topology of the HD space, and a gradient gene detector for on the fly retrieval of locally variable genes from the full gene set. We complement this with a suite of user-defined visualization options to inspect and query the data, and to project analysis results on the spatial data. We quantitatively validated SpaceWalker results on public datasets of varying data sizes, demonstrating that SpaceWalker accurately captures tissue architectural features, while at the same time retrieving locally expressed genes, in many cases coinciding with known marker genes. Finally, due to the localized computations on the kNN graph, Spacewalker scales very well to large datasets: only the pre-processing time increased (kNN graph computation), while on-the-fly exploration speed remained near constant for datasets ranging from 3k cells x 115 features to 170k cells x 483 features. Taken together, these results demonstrate that SpaceWalker enables linked visual exploration of tissue architecture and (spatial and HD) gene expression gradients, faithfully recapitulating known tissue architecture and biology in an interactive manner in large ST datasets.

### Limitations and Future Work

The use of distance metrics to compute neighborhoods in HD space in SpaceWalker may present a challenge known as curse of dimensionality, when the dimensionality of the datasets reaches a certain threshold. We tested SpaceWalker with a feature depth of maximum 483 genes, in Vizgen MERFISH data. Most results presented in this paper were based on cosine distances (manhattan distance combined with a shared distance metric22 based on overlap was used for HybISS data), because it presented a good balance between computational speed and robustness with respect to feature depth. In applications with larger feature depth (for instance where the spatial data is imputed with transcriptome-wide gene expression from sc-RNASeq data), the kNN graph could be constructed based on a PCA projection of the full data. The spatial gene filtering could then still be performed over the full imputed feature set, still enabling transcriptome-wide exploration of gradients over all genes.

At present, we implemented the peak filter by contrasting a small and a larger neighborhood (both in spatial and HD space). In this paper, neighborhood sizes for filtering were kept constant at a default setting. However, since the gene filtering is performed on the fly, the user can vary the filter sizes, where the ratio between inner and outer filter size shifts emphasis between peak detection and edge detection. The size of the filter in relation to the tissue structure can be adjusted accordingly.

As previously mentioned, Spacewalker can also be applied to data of the developing brain and revealed tissue structure related patterns. Though out of scope for this work, it would be interesting to further investigate SpaceWalker performance on multi-timepoint developmental datasets as well as extending SpaceWalker for exploring developmental trajectories. Moreover, while we have focused on the spatial map, the spatial filters can also be applied to any 2D maps, such as t-SNE and UMAP^29^ embeddings. For example, the spatial filters can be applied to a t-SNE map with a size approximately corresponding to the spatial size of subclass clusters, to identify subclass-specific genes, or expression gradients within or between clusters in t-SNE and UMAP scatterplots. Finally, in this paper we demonstrated that SpaceWalker can be applied to multi-slice ST data in two slices; Application to whole-brain ST atlases27 is currently under study.

Overall, SpaceWalker offers a novel and interactive visual interface for exploring ST data. By visually presenting the HD neighborhood structure of cells and genes with localized expression patterns, SpaceWalker helps study the tissue structure and identify potential genes of interest. SpaceWalker focuses on retrieving expression gradients and HD geometry on the fly, which differentiates it from the current state-of-the-art in script-based ST tooling, and is therefore complementary in its application and not a suggested replacement.

## Acknowledgments

This work was supported by the European Union’s Horizon 2020 research and innovation programme under the Marie Skłodowska-Curie grant agreement No. 860173; NWO Gravitation project: BRAINSCAPES: A Roadmap from Neurogenetics to Neurobiology (NWO: 024.004.012) and the NWO TTW project 3DOMICS (NWO: 17126).

## Author contributions

Algorithm development: C.L., J.T., T.H., B.L.

Data analysis: C.L., J.T., T.A., B.L.

Software Development: J.T., T.H.

Data interpretation: C.L., J.T., T.A., B.L.

Writing manuscript: C.L., J.T., T.A., T.H., B.L.

## Supplementary Figures

**Supplementary Figure 1:**
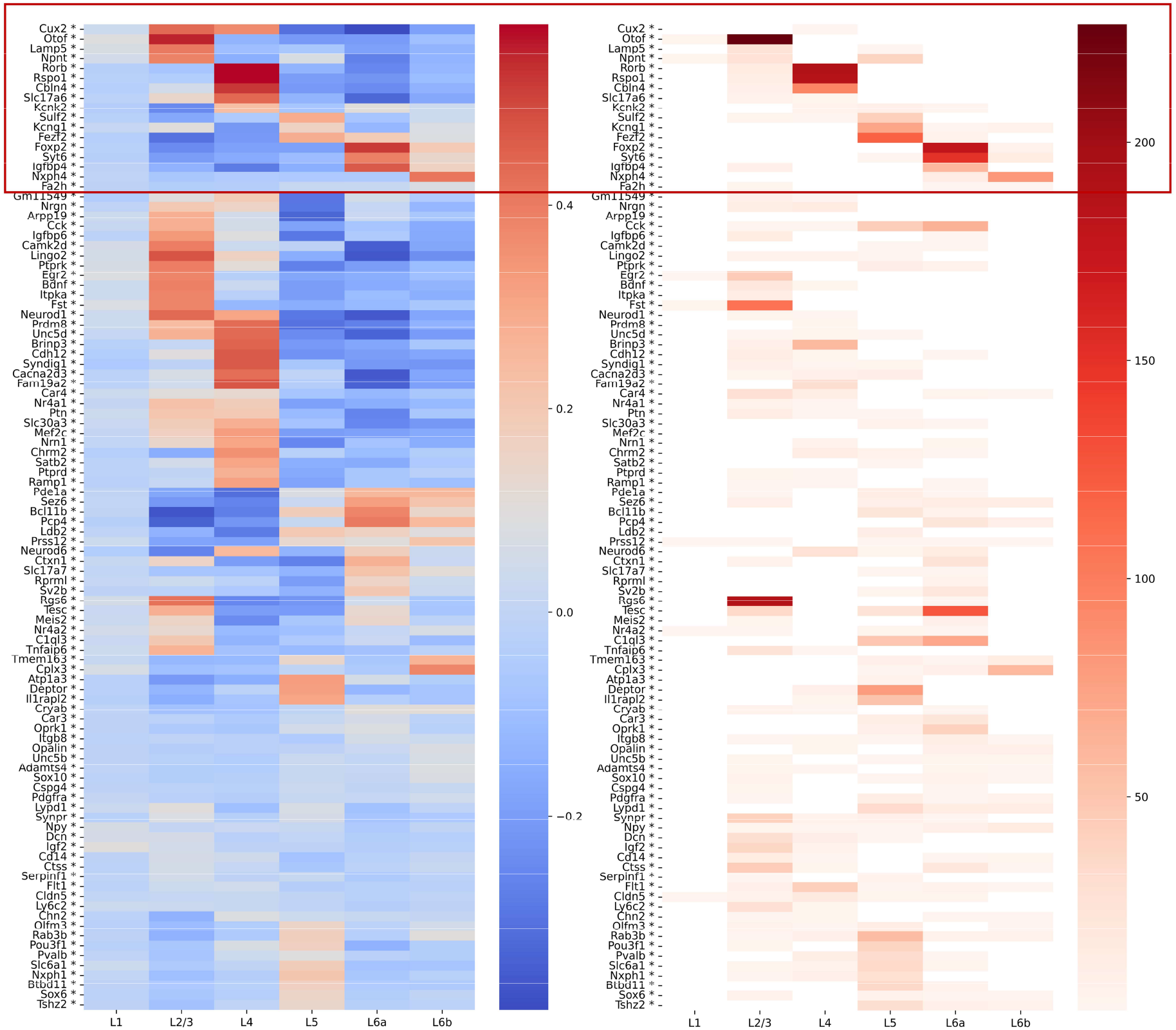
Heatmaps comparing the frequency counts by the spatial filter (right) with the correlation scores between genes and layer annotation masks (left) of MERFISH dataset. Known marker genes as reported25 are highlighted in the red box. Genes with a high correlation with a layer mask were also often ranked in the top two by the spatial filter.

**Supplementary Figure 2:**
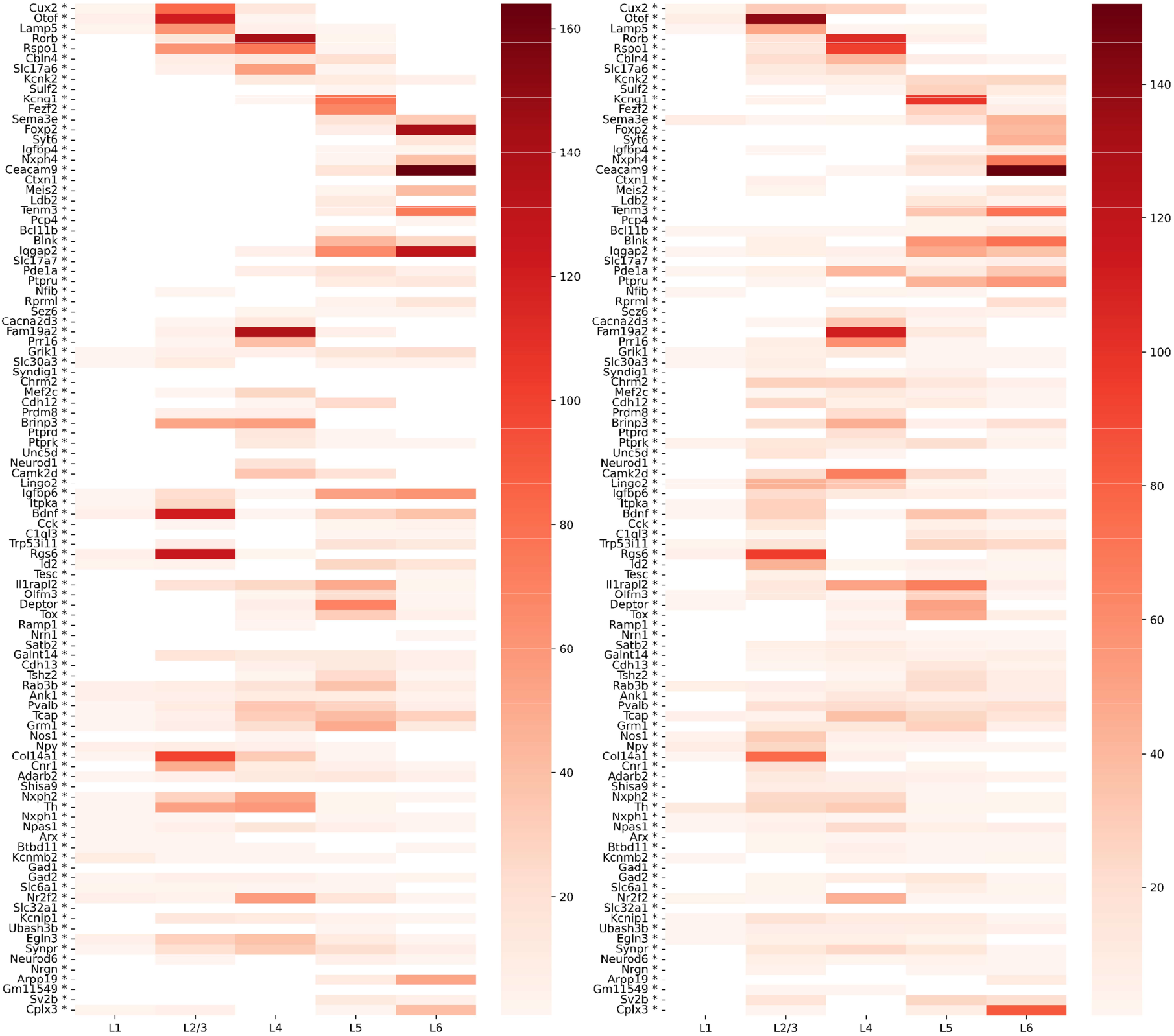
Heatmaps of the frequency counts by the HD filter in transcriptomics space (left) and heatmaps of the frequency counts by the localized spatial filter that only applied on flooded cells (right) of smFISH dataset. Similar to the results of the spatial filter (applied on all cells) in Figure 4, genes with a high correlation with a layer mask were also often top-two ranked by the HD filter and localized spatial filter.

**Supplementary Figure 3:**
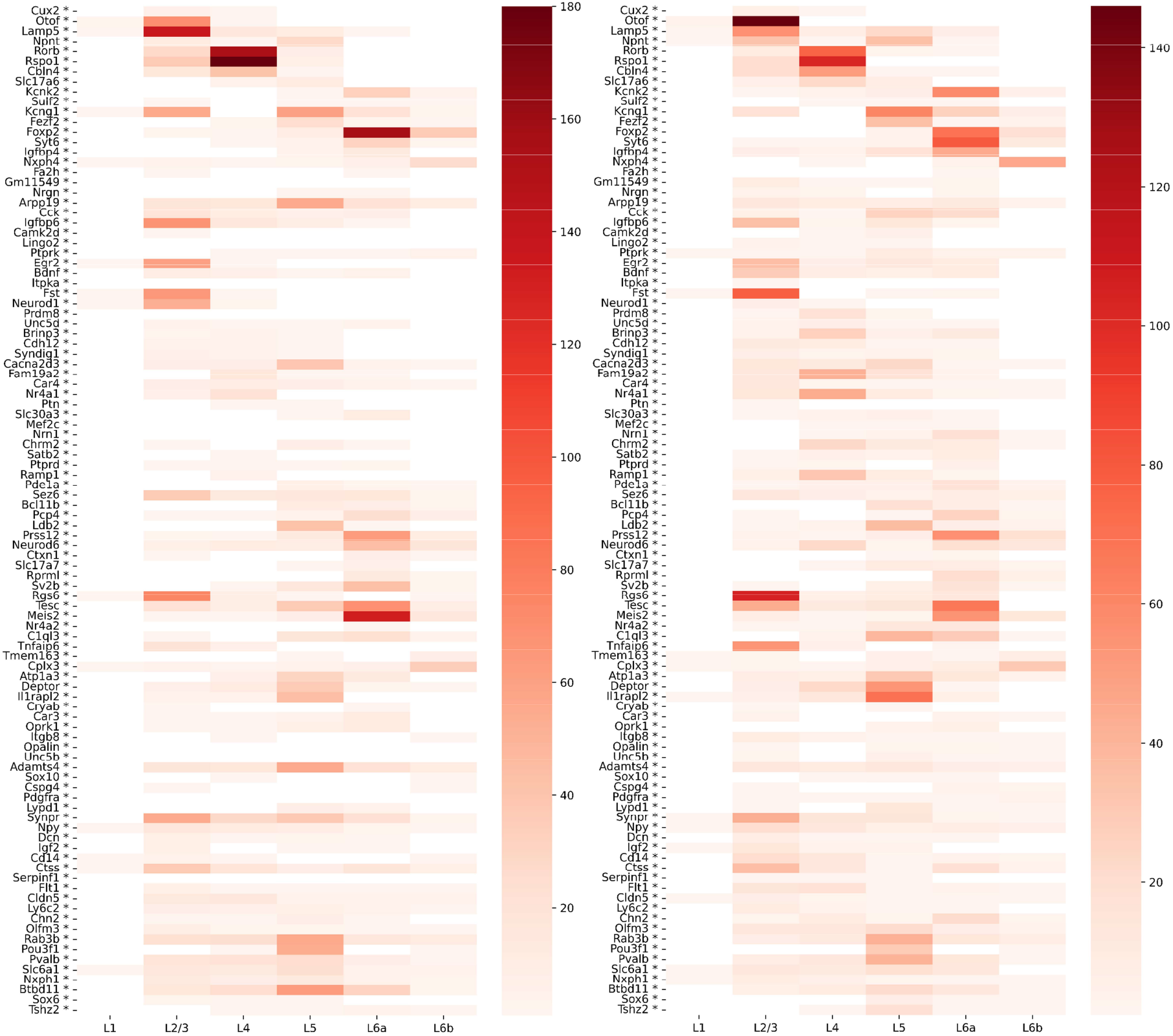
Heatmaps of the frequency counts by the HD filter in transcriptomics space (left) and heatmaps of the frequency counts by the localized spatial filter that only applied on flooded cells (right) of MERFISH dataset. Similar to the results of the spatial filter (applied on all cells) in Supplementary Figure 1, genes with a high correlation with a layer mask were also often top-two ranked by the HD filter and localized spatial filter.

**Supplementary Figure 4:**
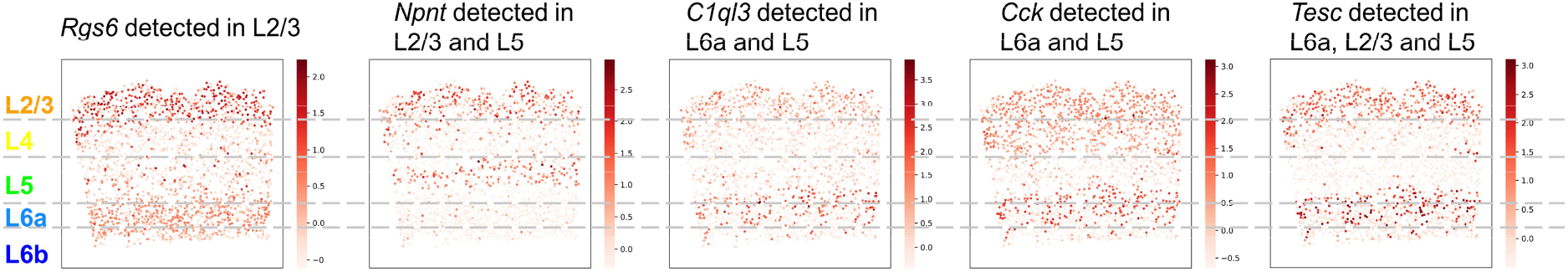
MERFISH spatial map color-coded by filter-detected genes that express in multiple layers. *Rgs6* is one of the most frequent genes that are ranked as top two in L2/3, and it also expresses in L6. *Npnt* is detected by the filter both in L2/3 and L5, and it expresses in both layers. *C1ql3* is detected by the filter in L5 and L6a, and it expresses in both layers. *Cck* is detected by the filter in L5 and L6a, and it expresses in both layers. *Tesc* is frequently ranked as top two in L6a, and it also expresses in L2/3 and L5.

**Supplementary Figure 5:**
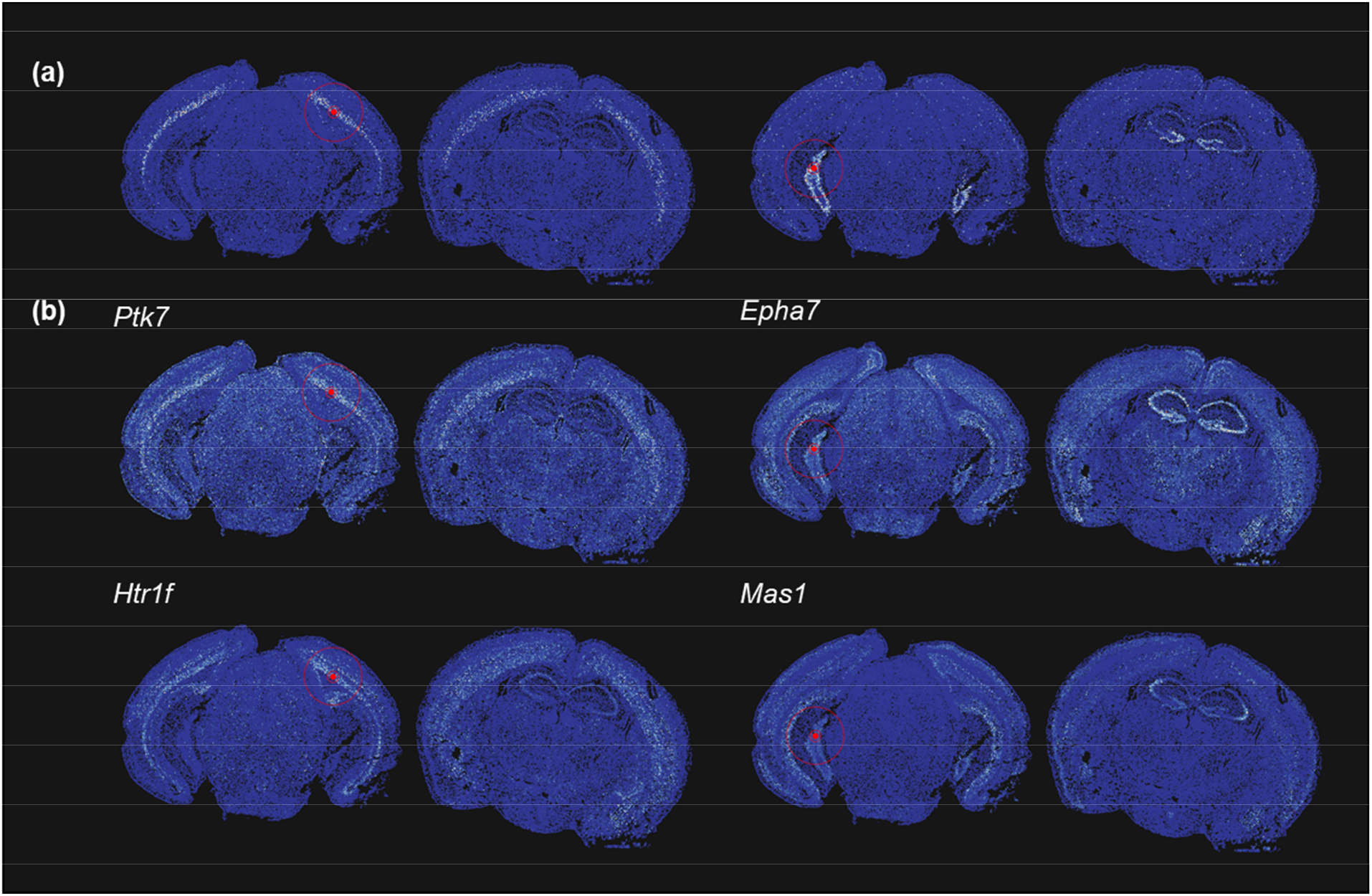
Results of multi-slice exploration of Vizgen MERFISH dataset. (a) Examples of local neighborhood reprojections, flooded cells are colored by flood-fill step index, highlighting similar tissue structure between slices. (b) Top two genes selected by spatial filtering at the location of the seed cell in (a), displaying localized expression patterns.

